# Meta plasticity and Continual Learning: Mechanisms subserving Brain Computer Interface Proficiency

**DOI:** 10.1101/2025.03.22.644643

**Authors:** Shuo-Yen Chueh, Yuanxin Chen, Narayan Subramanian, Benjamin Goolsby, Phillip Navarro, Karim Oweiss

## Abstract

**Objective:** Brain Computer Interfaces (BCIs) require substantial cognitive flexibility to optimize control performance across diverse settings. Remarkably, learning this control is rapid, suggesting it might be mediated by neuroplasticity mechanisms operating on very short time scales. However, these mechanisms remain far from understood. Here, we propose a meta plasticity model of BCI learning and skill consolidation at the single cell and population levels comprised of three elements: a) behavioral time scale synaptic plasticity (BTSP), b) intrinsic plasticity (IP) and c) synaptic scaling (SS) operating at time scales from seconds to minutes to hours and days. Notably, the model is able to explain *representational drift* – a frequent and widespread phenomenon observed in multiple brain areas that adversely affects BCI control and continued use.

**Approach:** We developed a closed loop, all optical approach to characterize IP, BTSP and SS with single cell resolution in cortical L2/3 of awake mice using fluorescent two photon (2P) GCaMP7s imaging and optogenetic stimulation of the soma targeted ChRmine_Kv2.1_. We further trained mice on a one-dimensional (1D) BCI control task and systematically characterized within session (seconds to minutes) learning as well as across sessions (days and weeks) with different neural ensembles.

**Main results:** We found that on the time scale of seconds, substantial BTSP could be induced and was associated with significant IP over minutes. Over the time scale of days and weeks, these changes could predict BCI control proficiency, suggesting that BTSP and IP might be complemented by SS to stabilize and consolidate BCI control.

**Significance:** Our results provide theoretical and early experimental support for an integrated meta plasticity model of continual BCI learning and skill consolidation. The model predictions may be used to design and calibrate neural decoders with complete autonomy while considering the temporal and spatial scales of plasticity mechanisms and their anticipated order of occurrence. With the power of modern-day machine learning (ML) and artificial Intelligence (AI), fully autonomous neural decoding and adaptation in BCIs might be achieved with minimal to no human intervention.

## MAIN

Implantable BCI technology have made striking advances over the last two decades, with numerous demonstrations of rapid and efficient neural control of prosthetic and communication devices by neurologically impaired subjects ^1–6^. At the heart of these advances are two important principles. The first principle is the neural decoder design which involves real time mapping of neural activity patterns to control signals that drive an agent in the task space– referred to as exogenous BCIs^7^ (e.g. a computer cursor or a multi degree of freedom (DOF) prosthetic limb^2,3^). It could also involve directly stimulating neurons based on decoded neural activity patterns to achieve a desirable functional outcome, such as inducing Hebbian plasticity between recorded and target brain regions^8^ (referred to as endogenous BCIs^7^). The second principle, which is a distinguishing feature of exogenous BCIs, involves the availability of sensory feedback once the BCI loop is closed; it allows the subject to learn which neural activity patterns are rewarded and hence should continue to be reinforced. It is believed that neuromodulatory signals play a key role in this process, both at the single cell level and the population level, allowing subjects to optimize BCI performance with repeated practice^9,10^ as in natural motor skill learning.

A notable observation in BCI studies, however, is the speed of learning, which can be in the order of a few minutes per degree of freedom^11–15^, although this depends on the training paradigm^16–18^ and the decoder design^19,20^. This is comparable to the quick, within-session motor skill learning but attaining BCI proficiency is remarkably faster than the slow learning process which takes many hours and days of repeated practice ^21^. Early attempts to explain these differences proposed that the standard Hebbian synaptic plasticity—resulting from (nearly) coincident depolarization of pre- and postsynaptic neurons—could be a biologically plausible mechanism that underlie this rapid learning. However, Hebbian plasticity occur on the time scale of a few tens of milliseconds^22–26^, which is insufficient to incorporate neurofeedback – an indispensable component in BCI learning^27^. Furthermore, co-activation of pre- and postsynaptic activity at larger intervals (seconds or longer) is known to produce no synaptic plasticity ^28,29^. More recent work suggested that the fast actions of dopamine D1 receptor (D1R) within 500 milliseconds was sufficient to drive reinforcement of the preceding cortical patterns^30^, which is consistent with the narrow time window (0.3-2 seconds) over which dopamine promotes dendritic spine enlargement following glutamatergic inputs ^31^.

Aside from plausible mechanisms of BCI learning, another notable observation in both BCI and non-BCI studies is the occurrence of *representational drift*– defined as a consistent shift in the representation of task variables despite that the associated behavioral and environmental conditions remain unchanged^32–38^. In BCI settings, this adversely affects performance and learning progress and necessitates frequent decoder calibration and possible corrective approaches to misclassified/missing data ^9,39–41^. Together, these observations and the lack of understanding of their underlying causes during continued BCI use have been a major impediment to BCI large scale translation with full autonomy^42 43^.

Herein, we propose a mechanistic model involving the integration of multiple distinct forms of plasticity spanning different time scales that are consistent with the above observations (Figure 1). The model comprises three elements: behavioral time scale synaptic plasticity (BTSP), intrinsic plasticity (IP) and synaptic scaling (SS). First, we posit that the primary mechanism for fast learning within the first few BCI trials is mediated by BTSP taking place over a few seconds ^44–47^, consistent with a neuromodulatory effect involving dopamine D1 receptor (D1R) activation within a sub-second timescale^30^. Second, we posit that the continued reinforcement of rewarded neural activity patterns triggers IP over relatively longer time scales (minutes). This process, in turn, results in perturbation of the homeostatic state of the circuits within which BCI neurons are embedded. The ensuing imbalance of excitation/inhibition (E/I) within these circuits triggers a reorganization process to restore the E/I balance through synaptic scaling^48,49^, a much slower process that operate over time scales of hours, days and possibly weeks. We posit that the interaction between these three elements reconciles multiple findings and provides a more accurate account for BCI learning, skill consolidation, and practical implementation during everyday use.

**Figure 1.**
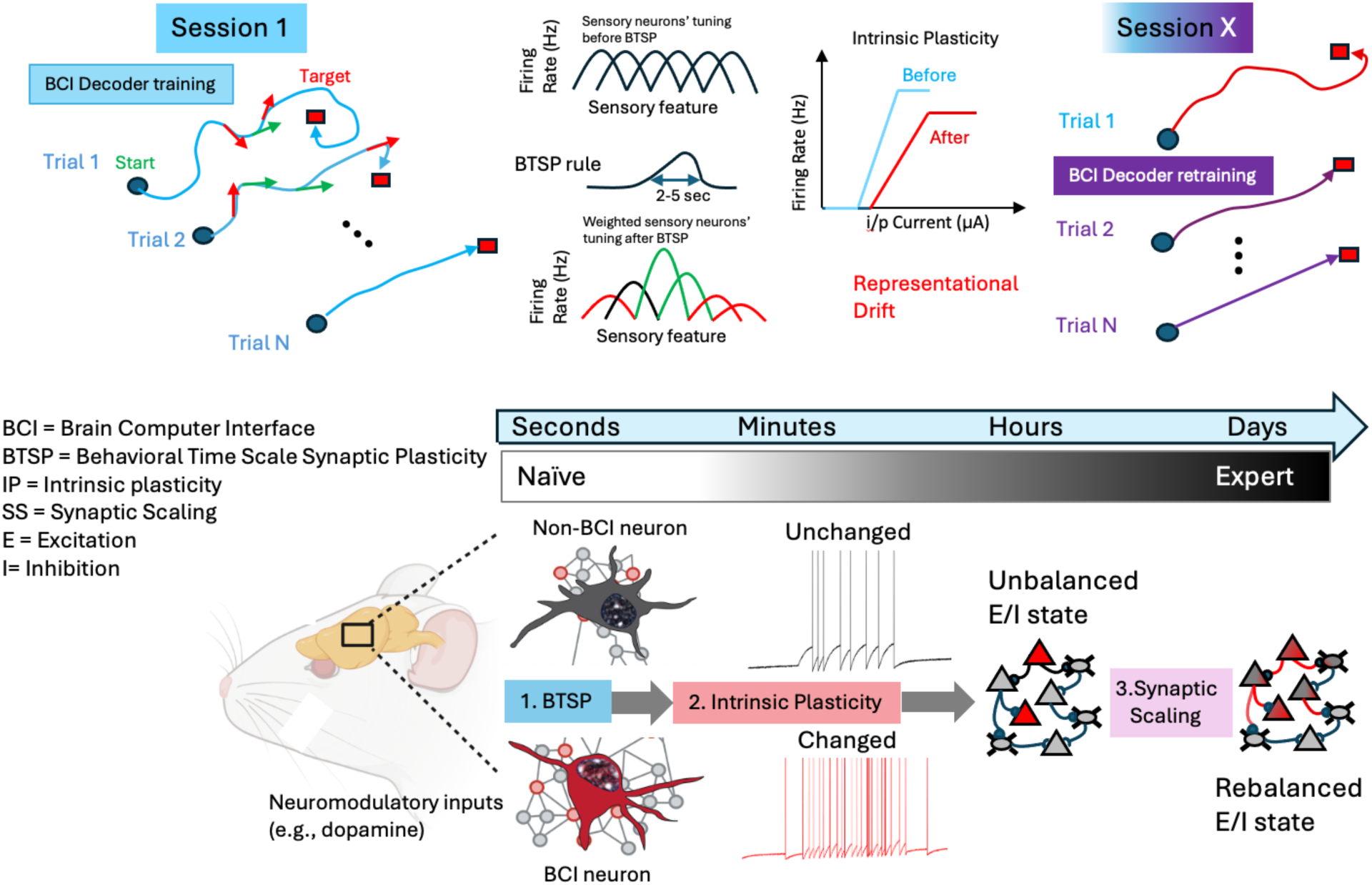
Meta plasticity model for BCI learning and skill consolidation featuring interaction between three elements: behavioral time scale synaptic plasticity (BTSP), intrinsic plasticity (IP) and synaptic scaling (SS) taking place over a multitude of time scales. During the first BCI training session (top left), rewarded neural activity patterns result in an increase in their frequency through BTSP (bottom left). The sensory feedback inputs arriving at different times along the cursor trajectory would then be potentiated if they are most relevant to the agent’s rewarded behavior (green arrows) and would be depressed if they lead to reward prediction error (red arrows). Through neuromodulatory influence and repeated activation, changes in intrinsic excitability result and may persist over longer periods, depending on the frequency of training (top and bottom middle). This shifted homeostatic state leads to an unbalanced excitation/inhibition (E/I) state that could also affect non-BCI neurons within the local circuits (bottom right). The balance is eventually restored through synaptic scaling over much slower time scales. The restoration of this balance might not re-instate the original excitability state of each individual neuron but may result in a new ‘attractor’ state of the ensemble representing the learned experience. These changes could explain the representational drift typically observed over similar time scales that adversely affects BCI performance and requires repeated BCI decoder calibration in subsequent sessions (top right).

### Rapid BCI learning and Behavioral Time Scale Synaptic Plasticity

BTSP is a short-term, non-Hebbian plasticity mechanism that is distinct from Long Term Potentiation (LTP), Long Term Depression (LTD) or Spike Timing Dependent Plasticity (STDP), in which presynaptic activity alters synaptic efficacies over behaviorally relevant time scale (seconds)^44,50^. It has been first demonstrated in hippocampal area CA1 neurons *in vivo* within a handful of trials during place cell formation ^44,50,51^. The basic idea is that presynaptic inputs that were neither causal nor close in time to postsynaptic activation can be potentiated if dendritic Ca^2+^ ‘plateau potentials’ are present around the dendritic structure where these inputs were received. These plateau potentials are generated by complex spiking and cause an increase in the weights of excitatory inputs from presynaptic neurons, essentially creating the so called ‘eligibility trace’ that interacts with an instructive signal (e.g. a reward input) to determine which synapses to potentiate. Most strikingly, the magnitude of this potentiation permits this phenomenon to occur without substantial repetition (1.4 trials under natural place field input, 5.1 trials under induced place fields in the hippocampus ^44^). A learning rule responsible for input potentiation would therefore span a much longer time (seconds) than predicted by standard Hebbian rules (tens of milliseconds) to include inputs that were not directly involved in driving neuronal firing. The dynamics of this process are strikingly similar to the time scale governing dendritic spine enlargement associated with near-coincident glutamate uncaging and dopaminergic activation ^31^.

It has been consistently observed that the agent control during BCI training relies crucially on sensory (mostly visual) feedback to make instant corrections to the BCI controlled variable(s). In the non-biomimetic approach to BCIs in which an arbitrary decoder is built based on the manifold directions of the BCI neural ensemble ^20^, neural activity has been characterized during the first few trials by complex spiking patterns ^12,13^, with variations spanning a few seconds, suggesting ‘a search’ could be taking place in the neural state space for an optimal neural spiking patterns to drive the decoder ^52^. This search would involve finding an optimal input to these neurons to evoke these patterns that maximize the decoder’s output in a statistical sense. It is therefore conceivable that, instead of multiple pre and post synaptic spike pairings within tens of milliseconds (as the standard Hebbian SP would require), an optimal input to BCI neurons would produce a single Ca^2+^ plateau potential in dendrites that last for tens of seconds ^47^. Dopamine release over a comparable time scale would therefore be sufficient to reinforce afferent sensory feedback information at selected dendritic spines, which would typically arrive hundreds of milliseconds later ^27,53,,54^. These inputs arriving at different times would then be potentiated if they are most relevant to the agent’s rewarded behavior (Figure 1). Specifically, the delayed, stochastic nature of these synaptic inputs would correspond to different error magnitudes in the task space, some would be small during trial epochs in which the agent’s behavior is aligned with the task goal (e.g. cursor trajectory points to the target and hence reward predicting^55,56^), while others would be large around epochs in which the agent’s behavior is not aligned with the task goal (e.g. cursor moving away from the target). As such, past synaptic inputs from co-active, presynaptic neurons and postsynaptic activation by dendritic plateaus facilitated by local dopamine release would alleviate the need for instant correlation of neurons controlling the BCI, and this would be sufficient to induce tuning of BCI neurons to the decoded variable(s) in very few trials consistent with a BTSP rule. This model predicts that a rapid transition from failed trials early in the training session to successful trials later in the session (i.e., fast learning) is a good predictor of an overall high performance both within the remainder of the session as well as across sessions.

### Representational Drift and BCI decoder calibration

While BTSP could explain the rapid and flexible control associated with early BCI learning, it cannot explain the need for frequent decoder calibration over multiple sessions ^9,19^. This calibration is intended to deal with uncontrolled changes in signal quality and information content that are typically observed over the slower time scales of days and weeks ^57^. These can be broadly categorized as: 1) degradation in signal quality that are associated with specific events such as neuronal displacement and glial reaction to device implantation which typically occurs within the hours and days following the event (but could persist over longer periods) ^58–60^ and, 2) shifts in information content in the form of changes in firing statistics despite that the associated behavioral and environmental conditions remain unchanged^33,38^. This representational drift has been observed in sensory and association areas of cortex as well as the hippocampus ^34–37^, and is not specific to electrical recording methods but have also been observed with optical imaging methods. It takes place at constant rates after a task has been learned and is substantially different from those associated with specific events ^32,33^. Most strikingly, drift rate in the same subject can differ across contexts, suggesting it may be a function of the behavior performed during the period of observation^61^. Recent computational modeling has proposed it could be tied to fluctuations in intrinsic excitability that could bias the reactivation of previously stored memories ^62^, or to activity-independent, stochastic synaptic processes ^63^. A notable observation is that, coincidentally, drift rates (days and weeks) seem to match the current timescale of BCI decoder calibration^60^, although this observation has not been systematically characterized.

Our second proposition is that as subjects learn to reinforce neural activity patterns during early BCI learning, BTSP-mediated changes in neural selectivity to the decoded variable(s) would be characterized by rapid reduction in variability in these activity patterns. During this early phase, neural activity essentially transitions from a highly irregular pattern to a highly regular and repeatable patterns with smaller variability during later intervals. This transition is desirable as it helps BCI source signals reduce variability necessary to stabilize the decoder output. However, it comes at a cost; maintaining these activity levels with reduced variability triggers intrinsic plasticity (IP) – an activity-dependent change in single neurons’ excitability in response to repeated depolarizing inputs.

IP has been widely documented in neocortex ^64^, cerebellum ^65,66^, hippocampus ^67^, among other areas ^68^. Despite being mostly characterized in brain slices *ex vivo* in which real physiologic signals such as motivation, attention, and reward are absent (but see ^69^), it has been postulated as a way for neurons to instantly adapt their spiking output to maintain their membrane voltage within a limited dynamic regime – known as a homeostatic state^70,71^. Neurons do so by changing a key structural parameter: the number of ion channel proteins present at the membrane, which occurs over a slow time scale of hours to days. It is believed that the cascade of these processes enables neurons to maintain stability and robustness of recently learned experiences and become more resilient to noise ^72–74^. However, this new network state and the ensuing spontaneous activity – in which cortex revisits specific neural patterns due to oscillations within cortical microcircuits ^75^ – locally drives further synaptic scaling^76^, and results in a re-organized neural circuit ^77^.

Our third proposition is that BTSP followed by IP alters the homeostatic state of the network in which BCI neurons are embedded and thus necessitates synaptic scaling ^48,71,76,78–81^. This is a process in which neurons re-adjust the strength of their synapses through glutamate receptor trafficking outside of behavior to stabilize firing rates ^79^. This alteration seems to be cell and synapse type ^82^, area and region specific ^79^, depending on the neural circuit connectivity and dynamics ^83,84^. The stochastic fluctuations of inputs to these neurons and altered synaptic efficacies outside of BCI training pave the way for representational drift to manifest in the form of altered spiking statistics, eventually requiring frequent decoder calibration ^19,85,86^.

## Methods

We first designed an endogenous BCI experiment to test the idea that both BTSP and IP can be triggered in persistently active neurons over behaviorally relevant time scales (seconds to minutes). We performed two photon (2p) Ca^2+^ imaging of Layer 2/3 excitatory V1 neurons in head-fixed mice during wakefulness, aroused state. Transgenic reporter mice (Ai162, TIT2L-GC6s-ICL-tTA2-D, JAX labs) were injected with a viral vector (AAV8-CamKIIa-ChRMine-mScarlett-kv2.1) to express the soma targeted light sensitive opsin, ChRmine_Kv2.1_^87^, in excitatory cell types expressing the genetically encoded Ca^2+^ indicator, GCaMP7s ^88^, followed by an implantation of a cranial glass window (4 mm diameter) over the injection site. After allowing 4-6 weeks for expression, animals were head-fixed under the 2P microscope (Bruker Ultima, WI) and the imaged Field of View (FOV, 900 μm x 900 μm) was screened for regions of interest (ROIs) between ∼100 and ∼250 μm below the cortical surface that co-express GCaMP7s and ChRmine_Kv2.1_. For each GCaMP7s^+^ and ChRmine_Kv2.1_^+^ neuron selected, we determined the lowest laser power (A_OL_) and optical pulse duration (PW_OL_) (*λ*=1040 nm) that depolarizes the soma, measured as a significant change in evoked Ca^2+^ Δf/f (*λ*=920 nm) relative to background activity (Supplementary methods).

To mimic the input needed for maintaining persistent, repeatable spiking activity patterns typically observed in BCI experiments, we designed a protocol for closed loop control of photo stimulation based on instant somatic Ca^2+^ readout to maintain the cell’s activity at a specific target level for a predefined interval (Figure 2). Intracellular somatic Ca^2+^ has been shown to be a reliable indicator of a cell’s spiking history, including dendritic plateau potentials^89,90^. This approach - referred to herein as an ‘optical clamp’^91^-permitted studying both transient and steady state response of individual cells to repeated depolarizing inputs *in vivo*. We then quantified the extent to which non-clamped neurons within the same FOV exhibited significant change in their activity dynamics as a result of clamping one or more neurons.

**Figure 2.**
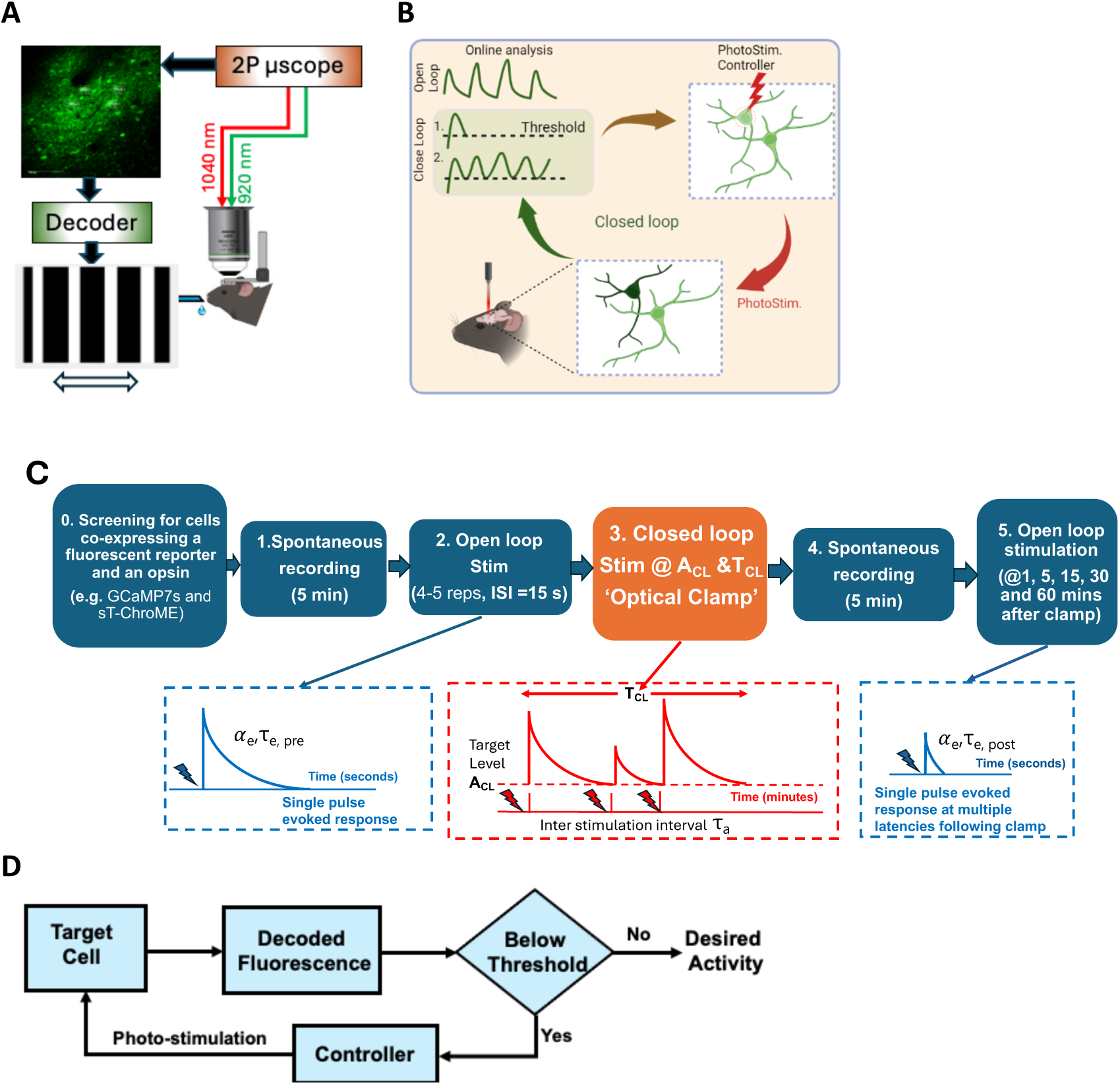
**A.** Schematic illustration of the exogenous BCI experiment. Rewarded neural ensemble activity instantly decoded and used as a control signal to drive a 1D ‘neural cursor’ on the screen in the form of orientation gratings. **B.** Endogenous BCI experiment uses the instantly decoded Ca^2+^ to directly photo stimulate the same (or different) neuron(s) co-expressing the opsin in closed loop at a non-overlapping spectral wavelength (1040 nm). The open loop photo stimulation is used to measure the effects of performing the closed loop approach at disjoint intervals. **C.** Protocol for characterizing BTSP and IP over a full experimental session. In the open loop step, each cell is photo stimulated with a single depolarizing pulse (4-5 repetitions, separated by 15 seconds for a total duration of ∼1 min) and the evoked Ca^2+^ traces are averaged to obtain a baseline for a proxy of the global dendritic plateau potential *before* the optical clamp. This step is repeated at multiple time points *following* cessation of the clamp. The parameters *α*_e_ and τ_e_ are used to characterize the changes in the plateau potential’s amplitude and time constant observed in each case. **D.** Endogenous BCI operation using an optical clamp: the decoded fluorescence is compared to a user specified threshold target activity level A_CL_, either in the same cell or in other cells within the FOV. If the activity drops below the target level, photo stimulation is delivered to depolarize the cell keep its activity above that level and the cycle continues for a user specified interval T_CL_.

### BCI control in vivo

We next designed an exogenous BCI experiment in which water restricted mice expressing GCaMP7s in excitatory neurons in layer 2/3 (L2/3) primary visual cortex (V1) were trained to volitionally modulate the activity of multiple ensembles of neurons for water rewards (Figure 2A). Given an ensemble, the animal was rewarded for modulating the average firing of a positive target pair (N+) above that of a negative target pair (N-) beyond a set threshold T1. Experiments were carried out over a number of sessions spanning multiple consecutive and non-consecutive days in which we alternated between ensembles that were either disjoint, partially or fully overlapping to quantify learning both within as well as across sessions (Supplementary methods).

## Results

We first tested whether single neurons could be optically clamped at different Ca^2+^ target levels for preset time intervals. After optimizing the optical stimulation parameters (pulse amplitude, pulse width and laser power) for each tested cell, we found that cells could be reliably maintained at target activity levels for many minutes following clamp onset (Figure 3A-B). We also found that multiple ‘target’ cells could be ‘yoked’ together with ‘trigger’ cells’ Ca^2+^ activity (Figure 3C-D), consistent with published reports ^92^. We also verified that off-target stimulation of adjacent cell bodies and/or neuropils was minimal to none (Figures 3E-3F).

**Figure 3.**
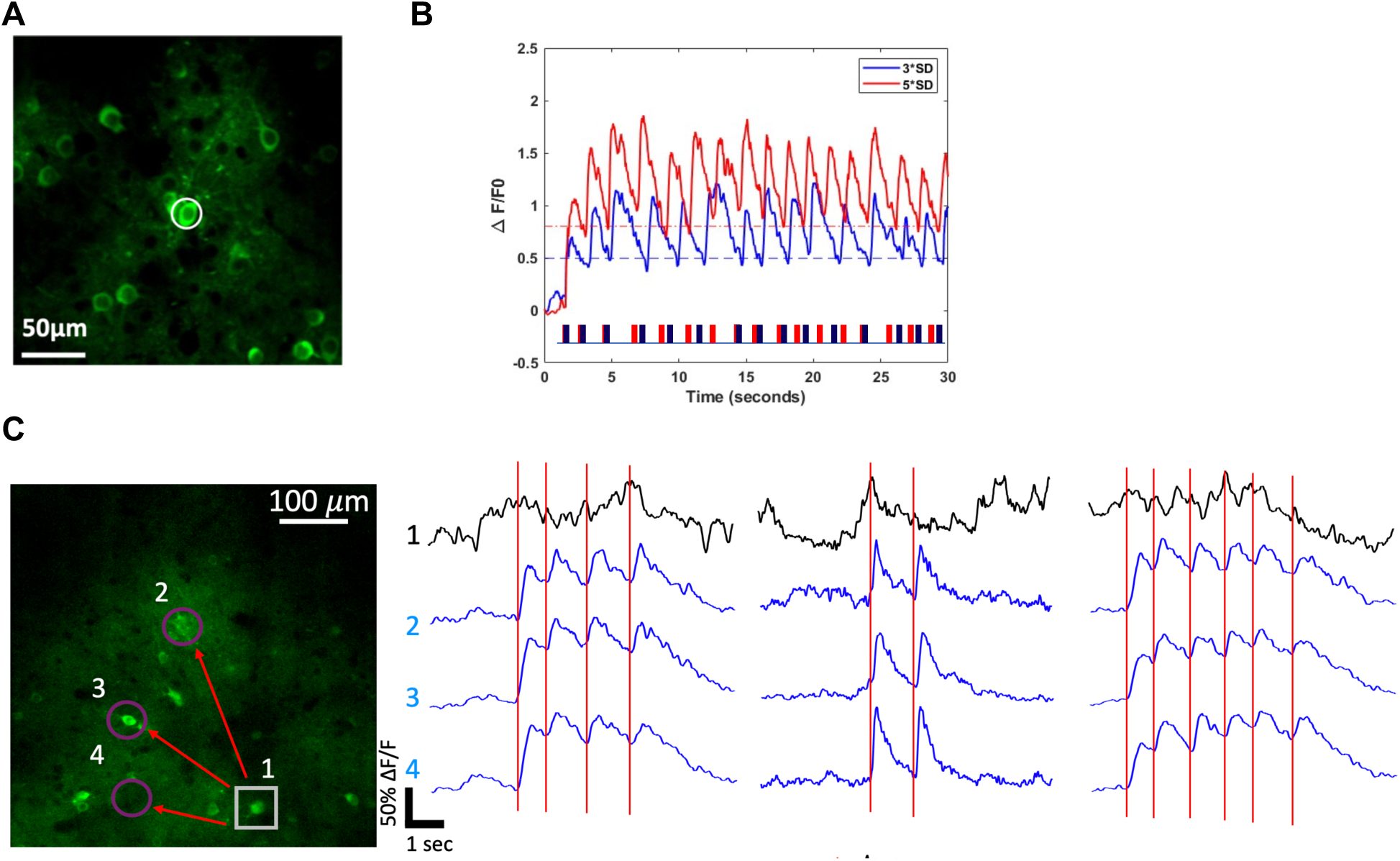
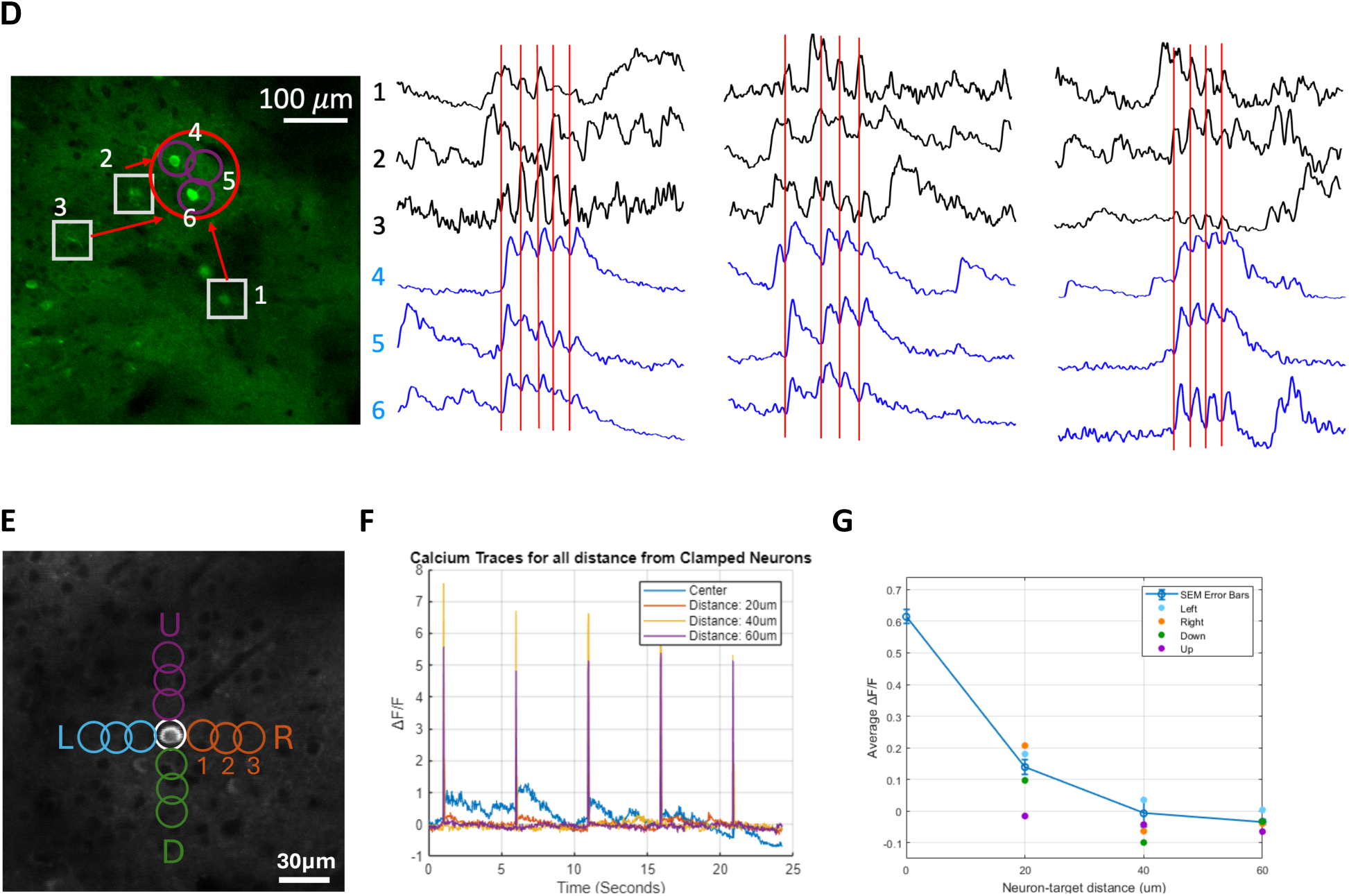
**A.** Example FOV with a single neuron (white circle) co-expressing ChRmine_Kv2.1_ and GCaMP7s targeted for closed loop photo stimulation based on the protocol in Figure 2B. **B.** Sample Ca^2+^ traces of the clamped neuron in A at two different target activity levels: mean + 3*Standard Deviation (SD) and mean + 5*SD for 30 sec. Ca^2+^ imaging was acquired at 30 Hz over an FOV of ∼250 × 250 μm using a resonant galvo scanning system at 920 nm and photo stimulated neurons via two-photon excitation at 1040 nm. The Spatial Light Modulator (SLM, Meadowlark Optics, CO) generated a single photo stimulation spot, which was scanned in a spiral fashion for 50 ms (5 spirals) per pulse. **C.** Examples of single and **D.** population optical clamping. Grey square and traces are *trigger* neurons. Blue circles and traces are stimulated *target* neurons that are ‘yoked’ together. Red lines indicate stimulation trigger times conditioned on activity decoded from the trigger neurons. **E.** A ∼250 × 250 μm FOV demonstrating the stimulation spots used to verify stimulation off target effects and 4 directions around the target cell body. **F.** Example Ca^2+^ traces from ‘off center’ color coded locations at 3 different spatial distances. **G.** Ca^2+^ fluorescence decay profile at photo stimulation spots ‘off center’ according to **E** and **F**.

We then asked whether the single cell optical clamp over behavioral time scales alters somatic Ca^2+^ signals in a way that mirrors the changes in the plateau potential. We reasoned that backpropagating action potentials (bAPs, which are ∼1 millisecond in duration^93^) evoked by depolarizing current injection into the soma result in broad subthreshold dendritic signals that are linearly associated with the number of action potentials produced at the soma, consistent with published work^45,94^. These bAPs are known to spread along proximal dendrites with noticeable attenuation at the farthest imaged locations^89^. Furthermore, following long depolarization, the somatic Ca^2+^ peak is expected to attenuate consistent with adaptation mechanisms associated with the reciprocal interaction between local dendritic spikes and depolarization-induced suppression of excitation^89^. Thus, we reasoned that somatic Ca^2+^ dynamics in response to a single depolarizing light pulse could be used as a proxy signal to estimate the changes associated with dendritic plateau potential following the optical clamp.

We stimulated each cell 4-5 times using a single pulse (50 ms, 5 spirals) separated by 15 seconds (total duration ∼ 1 min) and averaged the evoked Ca^2+^ traces to obtain a baseline for the global plateau potential before the optical clamp. We repeated the same step at 3 min and 8 min after cessation of the clamp. We found that when a cell is optically clamped for ∼200 sec (at 5*SD above its average baseline fluorescence), the evoked Ca^2+^ amplitude decreased by ∼18% (Figure 4A). Notably the decay time constant τ_e_ effectively decreased the response to ∼37% of its peak within 2257 ms ± 132.2 ms of the pre clamp window to 1850 ms ± 150.3 ms post clamp, consistent with BTSP models^44^. This signal on its own does not result in synaptic potentiation^95^. However, the overlap between this global signal and the local input signal at the dendritic compartment determines the degree of synaptic weight change for that input. Note that because our protocol relied on depolarizing the soma rather than the targeted activation of specific dendritic compartments, we could not measure the rise time of the plateau signal to verify its expected asymmetry ^44^. Nonetheless, the decay time produced in our model agrees closely with published studies.

**Figure 4:**
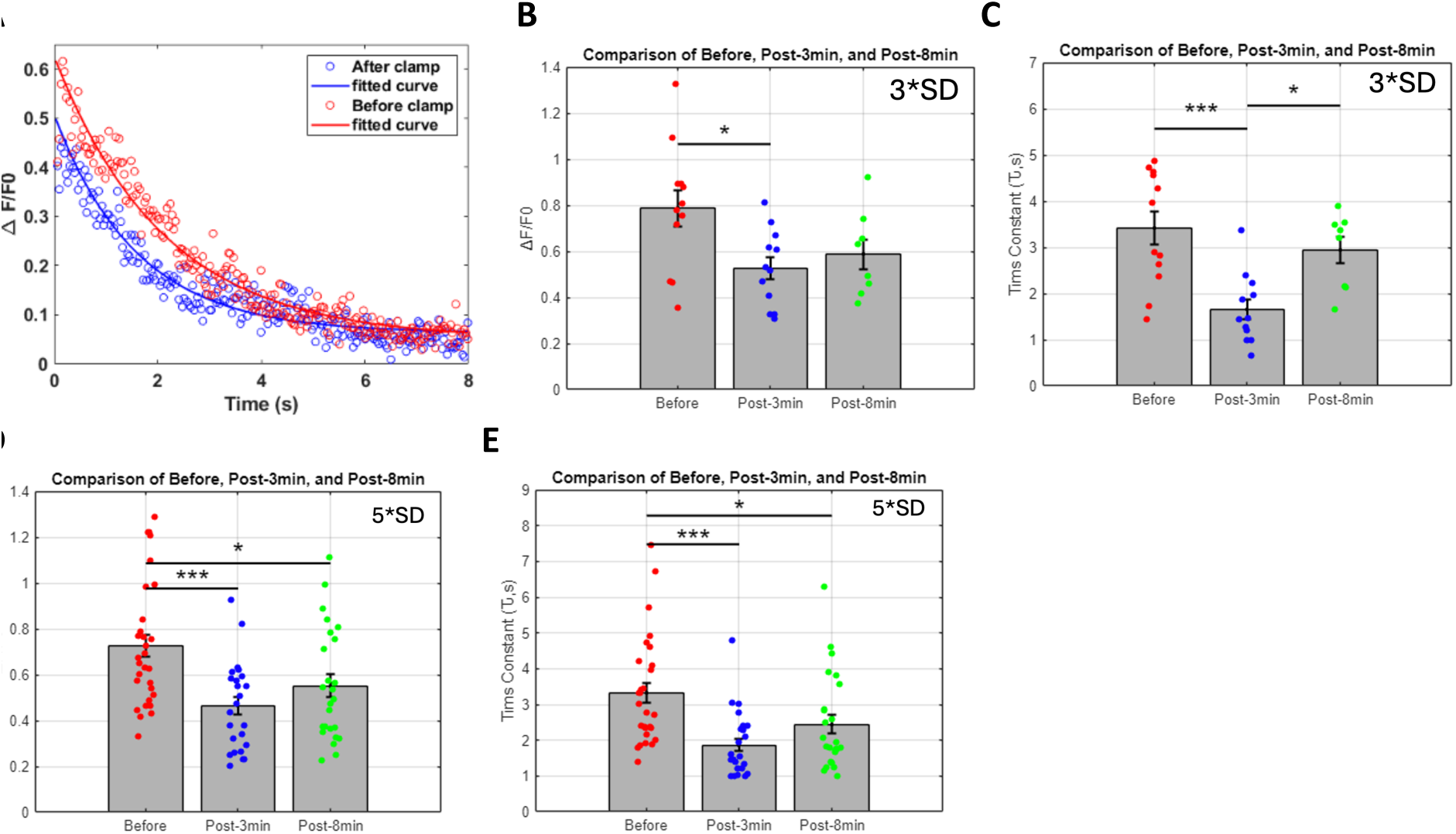
Characterization of somatic Ca^2+^ response to open loop stimulation. **A.** Example average change in Ca^2+^ impulse response to single pulse open loop stimulation, a proxy of the plateau potential, before and after optical clamp at 5*SD of baseline fluorescence for 200 seconds. **B.** Changes in peak Ca^2+^ amplitude *α*_e_ following clamp at 3*SD target level (before vs post-3min:*p=0.011035) that partially rebounded at 8 mins (before vs post-8min: p=0.065906; post-3min vs post-8min: p=0.47326) (n=3 cells, N=3 mice). **C.** Change in Ca^2+^ decay time constant τ_e_ following clamp at 3*SD target level (before vs post-3min:***p=0.00051785; before vs post-8min: p=0.31263; post-3min vs post-8min: *p=0.0032513) (n=3 cells, N=3 mice). **D.** Changes in peak Ca^2+^ amplitude *α*_e_ following clamp at 5*SD target level (before vs post-3min: ***p= 0.00010167; before vs post-8min: *p=0.015294; post-3min vs post-8min: p=0.1656). (n=7 cells, N=3 mice). **E.** Change in Ca^2+^ decay time constant τ_e_ following clamp at 5*SD target level (before vs post-3min: ***p=4.9042e-05; before vs post-8min: *p=0.024711; post-3min vs post-8min: p=0.076848). (n=7 cells, N=3 mice).

Across multiple experiments, we also found that the decrease in peak Ca^2+^ amplitude *α*_e_ at 3*SD target level was significant at 3 min (before vs post-3min: *p=0.011035) but partially rebounded at 8 mins (before vs post-8min:p=0.065906; post-3min vs post-8min: p=0.47326), suggesting the effect is transient (n=3 cells, N=3 mice; Figure 4B). The effect was more pronounced at 5*SD target level (before vs post-3min: ***p= 0.00010167) with slower rebound to pre-clamp levels (before vs post-8min: *p=0.015294; post-3min vs post-8min: p=0.1656) (n=7 cells, N=3 mice; Figure 4D). The changes observed in peak Ca2+ amplitude were mirrored by similar changes in decay time constant τ_e_. Specifically, we observed a significant decrease in τ_e_ at 3*SD target level (before vs post-3min: ***p=0.00051785; before vs post-8min: p=0.31263; post-3min vs post-8min: *p=0.0032513) (Figure 4C). At 5*SD target level, τ_e_ significantly decreased (before vs post-3min: ***p=4.9042e-05) and took longer to rebound (before vs post-8min: *p=0.024711; post-3min vs post-8min: p=0.076848; Figure 4E). These results suggest that Ca^2+^ dynamics are a function of the clamp target level and support the idea that perisomatic Ca^2+^ can serve as readout of the neuron’s activity in recent history. Furthermore, the observed effects are in general agreement with published work suggesting that after-hyperpolarizing current is depressed due to reduced postsynaptic CaMKII signaling ^96^. However, it is released gradually and reach baseline levels after one hour, a timing that coincides with the enrichment of several postsynaptic proteins to pre-plasticity levels^97^. These results suggest that a synaptic eligibility trace is generated that lasts for seconds to minutes following removal of a repeated depolarizing input.

We then asked to what extent the evoked Ca^2+^ response changes once the feedback control loop is closed, i.e., during optical clamp. We performed the optical clamp at multiple target activity levels (A_CL_ = 1*SD, 3*SD and 5*SD) for a preset interval (T_CL_ = 266 sec). Because the evoked Ca^+2^ response amplitude and decay time is measured during closed loop, characterizing the inter-stimulation interval (ISI) provides insight into how the cell adapts quickly to each new target. We consistently observed that the ISI was highly variable early during the clamp but later stabilized. To quantify these effects, we defined a *transient state* as the time interval from clamp onset during which the ISI variance is greater than 10% of its value at steady state. We expected that the larger the difference between the baseline activity level (before clamp) and the target activity level, the faster the response transition would be from the transient to the steady state dynamics. Simply stated, neurons are expected to adapt more rapidly (shorter decay time) to strong and persistent inputs, i.e. higher target level, compared to weak and sparse inputs. First, we found that the ISI variance was significantly higher in the first 60-sec compared to the subsequent second, third and fourth 60-sec intervals (Figure 5A). This trend was similar across all cells examined (n=5 cells, N=4 mice, Figure 5B). However, no significant differences were found between the second, third or fourth intervals for either the 3*SD or the 5 *SD cases. This trend persisted across all cells examined (Figure 5C-D). We also found that the ISI variance for 5*SD target level was significantly lower than the 3*SD level (Figure 5E), but there was no significant difference above 180 sec. These results suggest that: 1) the transition from early, largely irregular response to late, sustained and more regular response can be used to quantify how rapidly a cell adapts to specific target activity; the higher the target level the faster the adaptation and, 2) the time scale of this adaptation only lasts for 2-3 minutes. These transitions are likely cell dependent due to different morphologies and dendritic structures^89^.

**Figure 5:**
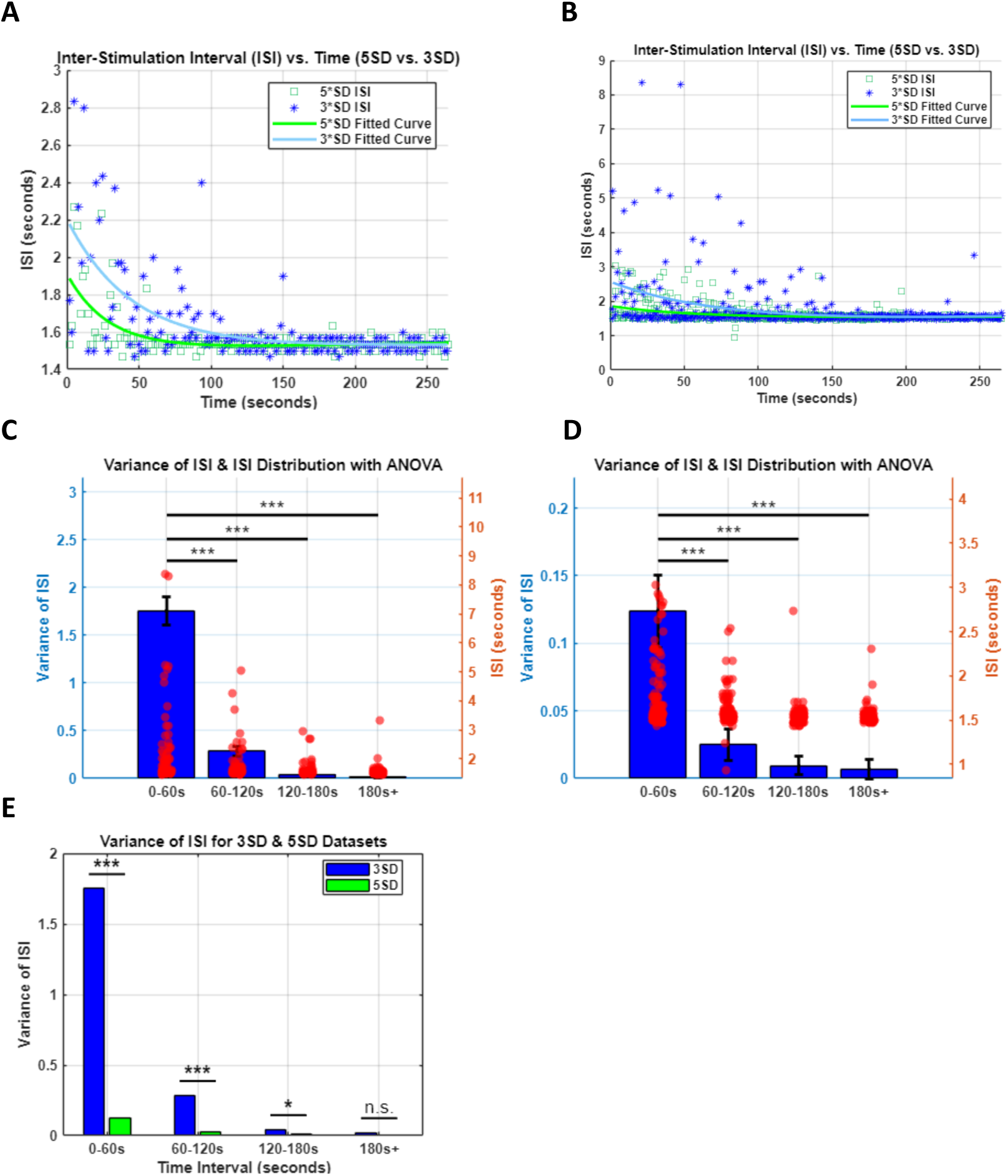
Characterization of somatic Ca^2+^ response to closed loop stimulation. **A.** Example Inter Stimulation Interval (ISI) and exponential curve fitting associated with clamping one example cell at 3*SD and 5*SD. **B.** Same as in A but for n=5 cells across N=4 mice. **C.** ISI and its variance during clamp broken down by 60 seconds intervals for target activity level 3*SD. Optical parameters used are power= 10mW; 161 pulses per 266s (or 36 pulses/min). 0-60sec, 60-120sec:***p=9.50E-06; 0-60sec, 120-180sec:***p=6.68E-10; 0-60sec-180+:***p=1.25E-12; 60-120sec, 120-180sec:p=0.385; 60-120,180+:p=0.132; 120-180sec,180+: p=0.964. **D.** Same as C but with target 5*SD resulting in 168 pulses per 266s (or 38 pulses/min). [0-60sec, 60-120sec]:***p=2.13E-09; [0-60sec, 120-180sec]:***p=2.36E-15;[0-60sec-180+]:***p=4.37E-13; [60-120sec, 120-180sec]:p=0.266; [60-120,180+]:p=0.350; [120-180sec,180+]: p=0.99. **E.** Comparison of ISI and its variance across two different levels across all cells in all mice (Two-Sample t-test: Time interval 0-60s: ***p = 0.00003; 60-120s: ***p= 0.00074; 120-180s: *p= 0.01464; 180s+: p = 0.64271)

We then asked whether the changes observed under both open and closed loop photo stimulation are associated with changes in the f-I characteristics. We performed 2P guided whole cell recording of select neurons and implemented the protocol in Figure 2. We then probed the cell’s input/output function by systematically varying the input power and measuring the spike rate (Figure 6A-B). We found that the power-frequency characteristics shifted significantly following clamp, suggesting response suppression. They further suggest that sub linearity at higher inputs takes place following periods of persistent activity. We then asked whether a computational model could capture these effects. We found that a Boltzmann function model with two parameters^98^, the adaptation time constant τ_a_ and the adaptation strength *α* can disentangle the relative contributions of each element of the feedback loop that relates the cell’s firing rate to the Ca^2+^ influx driven by the input current (Supplementary Figure 1).

**Figure 6:**
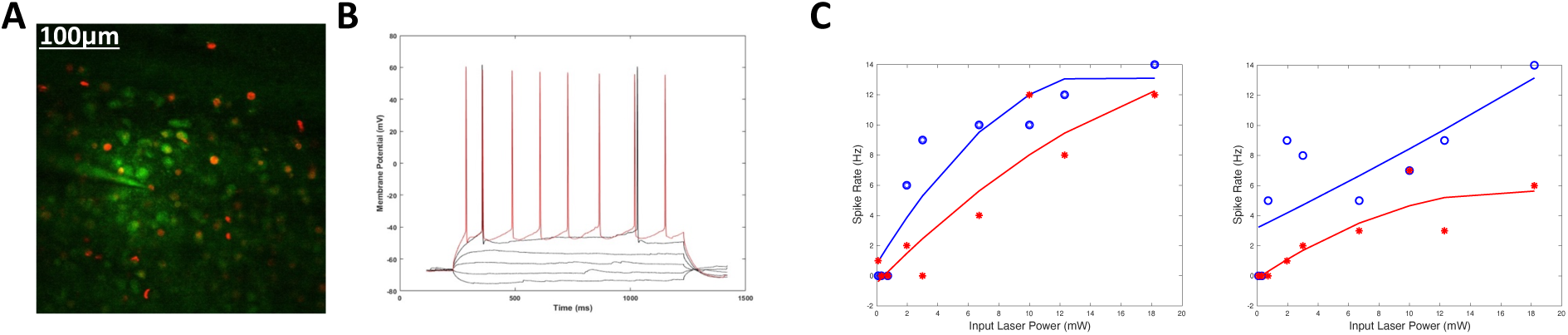
Characterization of changes in intrinsic excitability before and after optical clamp. **A.** Representative field of view for two photon-guided whole-cell recording in current clamp mode to characterize the cell’s f-I response. **B.** Sample whole cell membrane voltage recordings to two depolarizing current levels (5*μ*A black, 10 *μ*A red) and multiple non-depolarizing current levels. **C.** Power-frequency characteristics before and after clamp (at 5*SD target level for 200 sec) for two example cells. Blue open circles and red asterisks represent input laser power (proxy for the current injected) to spike rate before and after clamp, respectively.

Recent studies have suggested that neurons within Layer 2/3 of the primary visual cortex (V1) with similar stimulus feature selectivity are more likely to be synaptically connected compared to cells that have dissimilar tuning ^99^. We therefore asked whether clamping one cell’s activity at a particular target level has any influence on other adjacent cells within the same FOV. We clamped one cell at different target levels and quantified the changes in Ca^2+^ in the other imaged ROIs in terms of peak fluorescence and decay time constant during clamp. We found a significant increase in Ca^2+^ fluorescence intensity and decay time constant in n=17/46 cells response to target level (5*SD) (p = 0.0011; two-sided t-test), and for target level (3*SD) n=9/46; p= 0.0045; two-sided t-test) (Figure 7, Supplementary figure 2). While our experimental design did not permit mapping synaptic connectivity between the imaged neurons, published reports ^99^ suggests that optically triggering action potentials in a targeted neuron and directly measuring its functional influence on neighboring, non-targeted neurons can be used to assess synaptic connectivity profiles and accordingly make predictions about the extent of synaptic scaling taking place over longer periods.

**Figure 7.**
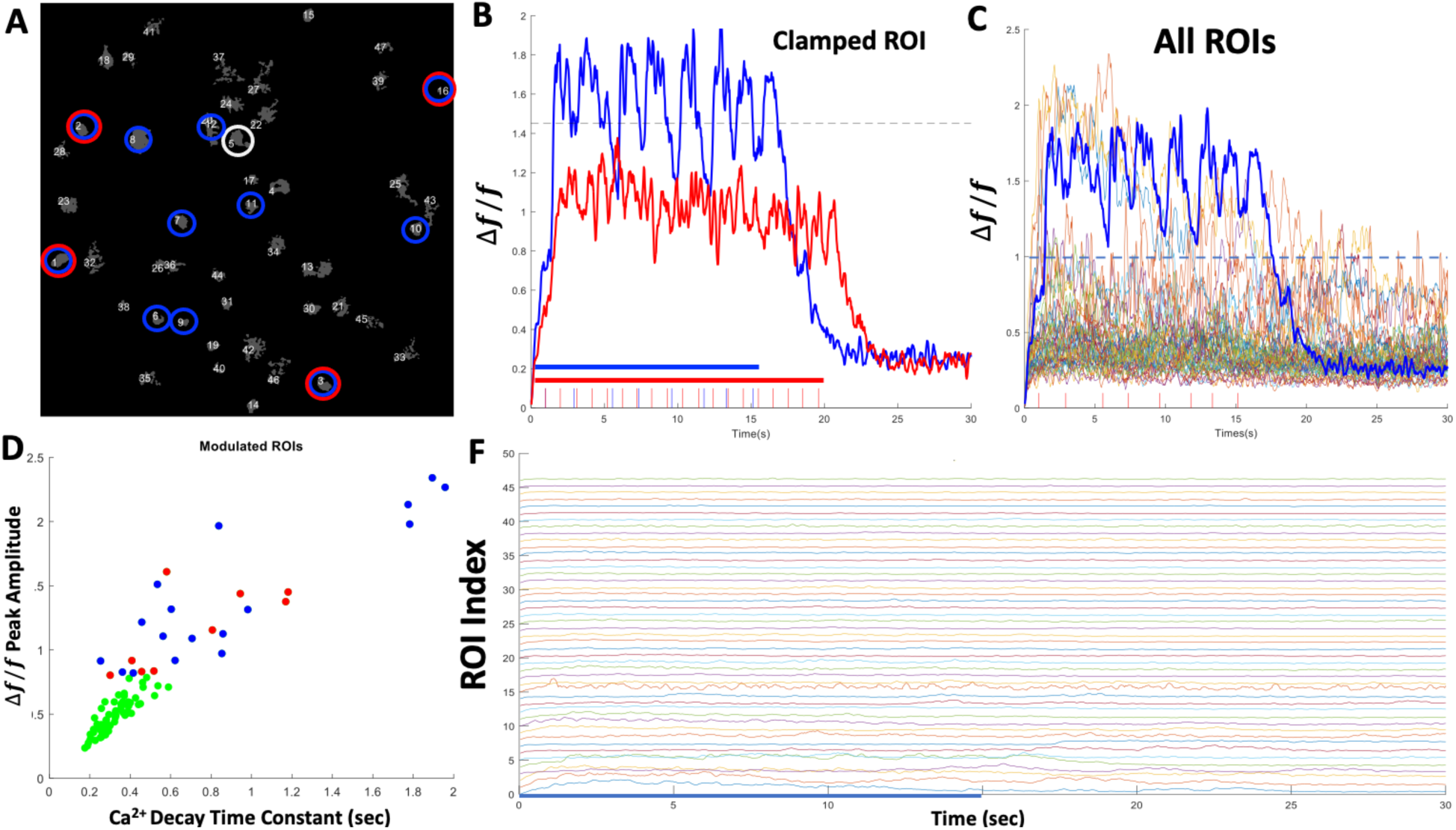
Influence of clamped neuron on adjacent neurons within the same FOV. **A.** Example FOV containing 46 well isolated ROIs in mouse V1. ROI # 5 (circled in white) was selected for optical clamping at two target levels shown in B. ROIs circled in red are cell bodies showing significant increase in Ca^2+^ fluorescence intensity in response to target level (3*SD) in B. ROIs circled in blue are cell bodies showing significant increase in Ca^2+^ fluorescence intensity in response to clamp level (5*SD). **B.** Activity of the clamped ROI#5 in A and photo stimulation pulse trains used for each target level (red 3*SD, blue 5*SD). **C.** Raw Ca^2+^ fluorescence of all ROIs in the FOV during clamp interval for target 5*SD for comparison. ROIs showing significant modulation synchronized with the clamped ROI (n=17/46 cells, 5*SD target, **p = 0.0011) correspond to blue circled ROIs in A. **D.** Ca^2+^ peak fluorescence versus decay time constant for all 46 ROIs in the FOV for target level 5*SD in C. Each dot represents the average of Ca^2+^ peak fluorescence and event decay time constant locked to each photo stimulation pulse during the clamp. Green dots are ROIs that were not significantly modulated within the clamp period (n=29; p=0.152). **F.** Raw Ca^2+^ fluorescence for all 46 ROIs in A at target 5*SD. Significantly modulated ROIs are plotted consecutively to demonstrate the difference between the activated and non-activated ROIs shown in C following cessation of the clamp at 15 sec.

Finally, we asked whether the above findings carry over to actual behavioral experiments of BCI learning and skill consolidation. We trained mice on a 1D BCI control over many weeks (supplementary methods). In six separate sessions, we selected six different ensembles of four neurons each (Figure 8A) and used a decoder filter to calculate the position of the cursor (orientation gratings moving at a fixed angle) at each time step. The gratings bars increased in thickness closer to the target and vice versa. Following an initial random cursor location at the start of each trial, the animal was rewarded if neuron pair N+=(N1& N2) increases the firing rates while neuron pair N-=(N3, N4) decreases the firing rates until the cursor attains the target (all black screen). Thus, the animal had different strategies to attain a target; for example, it could activate either or both neurons of the group N+ or decrease the activity of neurons in the group N-. Moreover, even if N+ neurons increase their activity, the activity of neurons in N-may be elevated enough to prevent the neural cursor from reaching the target. As a result, not all ensemble neurons were active at the time of target acquisition and not all activation of ensemble neurons resulted in a target.

**Figure 8.**
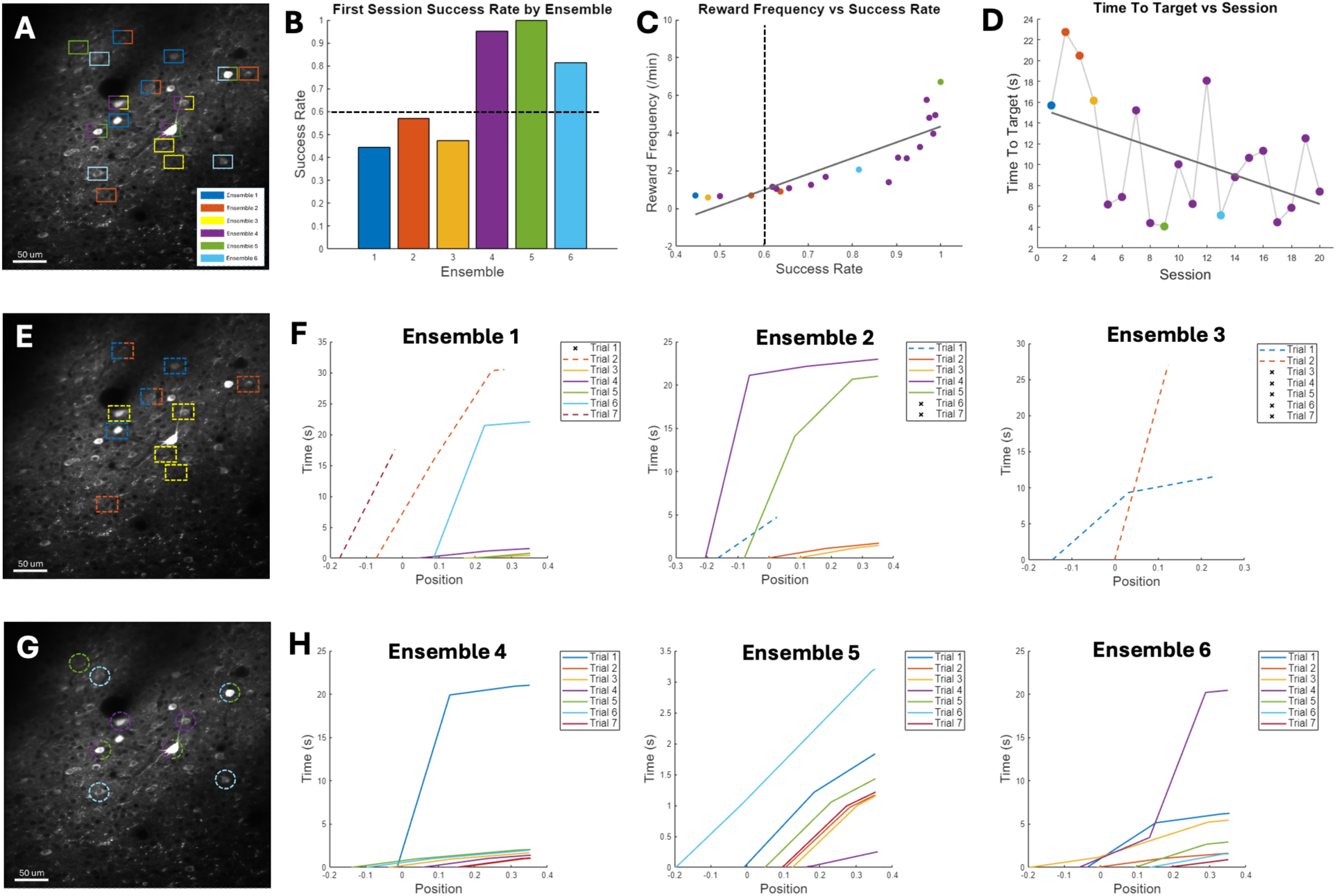
**A.** FOV where six different ensembles of 4 neurons each were used to train the animal on the BCI task over 6 different sessions. Some ensembles overlapped whereas others did not. **B.** Success rate in the first session for each of the ensembles in **A**. Dashed line indicate a threshold of 60% success that was used to categorize the ensembles into either a ‘weak/slow learner’ (E1-E3) or a ‘strong/rapid learner” ensemble (E4-E6). **C.** Average Time to Target in each session (color coded by ensemble) demonstrating continuous decrease over sessions for ensemble E4. **D.** Time to Target (TT) over sessions for ensemble E4 and for session 1 of the other 5 ensembles. Strong learner ensembles E4-E6 exhibited a TT= 6.21± 6.1 sec over 20 sessions compared to TT= 17.8 ± 20.2 sec for the weak learner ensembles E1-E3. **E.** Same FOV as A but highlighting only the neurons selected for ensembles E1-E3 using the same color code in A. **F.** Cursor path from initial start position to target position in the first 7 trials of the first session for ensembles E1-E3. Dashed lines indicate incomplete trial, whereas x indicates a failed trial (timeout). **G.** Same FOV as A but highlighting only the neurons selected for ensembles E1-E3 using the same color code in A. **H.** Cursor path from initial start position to target position in the first 7 trials of the first session for ensembles E4-E6.

We characterized the animal’s learning over two weeks of training and found that the animal’s increased reward frequency and decreased time to target (TT) (Figure 8B, 8C and 8D). We then examined the animal’s behavior within the first training session for each ensemble. Owing to the causal mapping between the BCI neurons and cursor movement, we used a simple behavioral criterion to compare the success rate in the first session for each ensemble to measure the extent to which rapid learning occurs within that ensemble. Our model predicts that re-occurrence of specific spiking patterns would be facilitated by BTSP taking place within a few trials. We set a criterion of 60% success rate to separate weak/slow from strong/rapid learner ensembles. Among the six ensembles investigated (n=24 total BCI cells, Figure 8A), we found that three ensembles were categorized as weak/slow learners in which within session success rate was ∼49% (Figure 8B, 8E and 8F), with significantly long Time to Target (TT) (n=12, ***p=0.002). In contrast, the other three ensembles had a success rate of ∼92%, with significantly smaller TT (Figure 8B, 8G and 8H) (n=12, ***p=0.0001). These results suggest that learning might not progress at the same rate in every ensemble, and that the role that each neuron plays within an ensemble may differ across ensembles.

Our model predicts that with repeated activation of an ensemble, changes in homeostatic state can be measured as changes in baseline firing rates across sessions. We selected one of the strong/rapid learner ensembles (ensemble E4) and continued to train the animal on the BCI task for an additional 13 sessions spanning a two-week period. We then compared the firing rate of BCI neurons during behavioral trials. We found no significant change in the N+ group (n=12; two sample t-test; N1: p=0.18, N2: p=0.07) between the first session and the last session (Figure 9A, Supp Fig 3A). These results suggest that the animal did not change its strategy with respect to modulation of the N+ group. In contrast, there was a significant change in the N-group (N3: ***p= 5.6e-10, N4: ***p=2.05e-05). Notably, these changes were opposite to one another; BCI neuron N3 decreased its firing whereas BCI neuron N4 increased its firing such that the net result among the N-pair was negligible, resulting in similar performance (90% in session 1 and 98% in the last session). We also found no significant changes in non-BCI neurons’ spiking probability during behavior (Supplementary Figures 3). These results suggest that the animal changed its strategy for modulating the N-pair between the two sessions that were examined albeit this change had no net effect on behavior.

**Figure 9:**
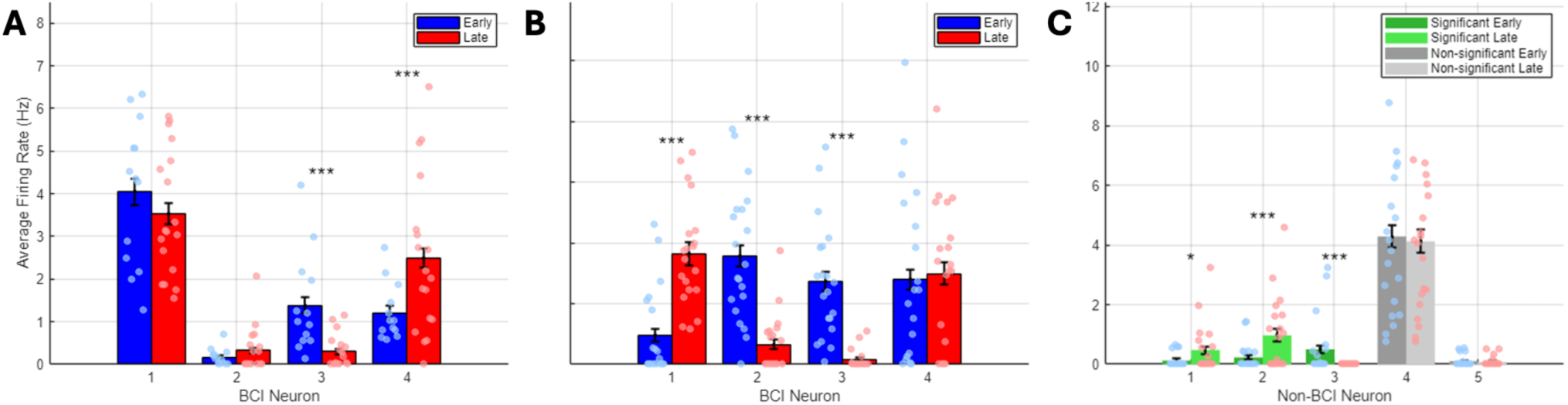
**A.** Average firing rate of BCI neurons during BCI control in the same ensemble (E4) comparing an early session versus a later session (2 weeks apart). Each dot represents the average firing rate over a 50 second interval within ∼10 mins of behavioural trials. Success rate in both sessions was 90% and 98%, respectively. **B.** Baseline firing rate comparison for the same ensemble in **A** outside of BCI behavioural sessions. Each dot represents the average firing rate over a 50 second interval within ∼8 mins. **C.** Baseline firing rate of non-BCI neurons in the same sessions as A. Each dot represents the average firing rate over a 50 sec interval within ∼8 mins (p=0.013;0.00085;3.77e-05;0.756;0.598 for neurons N1-N5 respectively, two-sided t-test)

To further examine changes in homeostatic states, we compared spiking probabilities and firing rate (outside the task) of neurons within the same ensemble around the early and late sessions (supplementary methods). We found significant changes in BCI neurons N1-N3 (***p=8.6e-10, ***p=7.72e-13, ****p= 5.75e-15) but not for N4 (p=0.68) (Fig 9B). We also found similar trends in non-BCI neurons within the same FOV (Fig 9C, supplementary Figure 4). These results suggest that some BCI neurons recruited during learning undergo changes in homeostatic firing, depending on the strategy used by the animal to optimize control. These changes could also affect non-BCI neurons within the local circuit as our analysis of non-clamped neurons within the same local area has shown (Figure 7; supplementary figure 2).

## Discussion

In this study, we proposed an integrated plasticity model for BCI learning that might explain multiple observations widely documented but have not received sufficient attention in the BCI community. This meta plasticity model, comprising synergistic interaction between multiple forms of neuroplasticity, could account for rapid within session learning, representational drift over days and weeks, and BCI skill consolidation. We developed a novel protocol to probe BTSP and IP that relies on targeted closed loop all optical control of single cell activity dynamics and showed that, on the time scale of seconds to minutes, substantial BTSP could be induced. The ensuing changes in neuronal firing statistics are consistent with known reciprocal interaction between local dendritic spikes and depolarization-induced suppression of excitation.

We first showed that open loop photo stimulation before and after perisomatic Ca^2+^ perturbation events such as an optical clamp can be used to measure global (but not local) plateau potentials. The perisomatic region of pyramidal cells is defined as a domain of plasma membrane that receives almost exclusively GABAergic synapses, which includes the soma, the axon initial segment, and the proximal apical and basal dendrites up to a distance of ∼100 μm. These dendritic compartments contain only very few spines and receive hardly any excitatory synapses ^100^. This direct photo stimulation approach to the cell body therefore bypassed possible perisomatic inhibition that might be triggered by recurrent inhibitory connectivity.

We then showed that following artificial induction of the same activity pattern in single cells using an optical clamp, significant adaptation occurs over 2-3 minutes in response to repeated depolarizing inputs. While this was not surprising given prior reports, we found that this adaptation is a function of the target activity level. These results suggest that maintaining these activity levels rapidly sculpts neurons’ dynamics to produce a delimited alteration of firing consistent with the input pattern. This feature could endow single neurons with the capacity for rapid memory formation representing that input pattern. We then showed that over the time scale of minutes following cessation of the clamp, rebounding of perisomatic Ca^2+^ homeostasis takes place. These results are consistent with the behavior of hyperpolarizing currents that are released gradually and reach baseline levels around the timing of enrichment of several postsynaptic proteins to pre-plasticity levels.

For neurons to engage BTSP, they need to act as a feedback control module that continuously monitors their own firing output in response to a particular input pattern and generates a spatially distributed synaptic eligibility trace consistent with that input via a Ca^2+^ mediated plateau potential. The neuron would then potentiate the inputs corresponding to that trace when they contribute to rewarded actions over seconds long interval^71^. This time window is critical as it permits the ‘time to reward’ to be simultaneously encoded alongside the eligibility trace that might last for a few trials^101^. Indeed, prior work using Ca^2+^ imaging in cortical association areas has shown that neurons continue to encode past trial experience^102,103^, including reward omissions, suggesting that these eligibility traces are critical for action-outcome history. If the neuron’s output is within the ‘target’ level, this global instruction signal should decay relatively fast, as there is no more ‘errors’ to correct. If the neuron’s output is significantly deviating away from that target level, the instruction signal should last longer to potentiate possible inputs needed to correct these errors, consistent with reward prediction errors of dopaminergic modulation.

To bypass perisomatic inhibition that might be caused by recurrent connectivity and further decouple the role of reward and associated dopaminergic influence on the induced plasticity, we designed an experiment to characterize the BTSP mechanism by mimicking the recurring activity pattern observed in BCI neurons– an elevated excitability state – in a closed loop control setting. To simplify the analysis and to avoid potential confounding factors, we simplified these activity patterns to an average target activity level that the neurons needed to maintain above their baseline for a predefined time interval. It is important to note that our goal was not to mimic the full repertoire of activity patterns typically observed during volitional modulation of BCI neurons, but rather to study how different target activity levels that neurons must maintain could result in significant changes in perisomatic Ca^2+^. We believe this design provided insights into how these signals could be critical in representing the instructive signals in the early few trials during learning.

We then provided preliminary evidence supporting the idea that the altered spiking probabilities could trigger changes in the intrinsic excitability of the neurons. We have shown that re-occurrence of activity patterns close to the selected target induces IP and shifts the f-I characteristics towards response attenuation and possibly sub linearities. Our modeling results further recapitulated this effect, lending credence to our model structure. We then showed that neurons within the ‘broader circuit’ may also be influenced by the transient and sustained changes in the homeostatic state of the targeted neurons, despite not being directly targeted by photo stimulation. While it was not feasible to map synaptic connectivity between neurons within the local population, we showed that a substantial fraction of the population within the local area can exhibit significant changes in their activity dynamics. Modern approaches, including our own work ^104,105^, to use two photon optogenetics to map synaptic connectivity could play a crucial role in this regard.

Several reports have used recurrent neural network (RNN) models equipped with a reward modulated Hebbian plasticity mechanism to demonstrate that neuromodulators such as dopamine could elongate the time window of integration between efferent and afferent BCI signals to enable learning simple BCI tasks ^106–108^. However, with the exception of a recent study^30^, these models lack experimental validation at the cellular level and have focused on average learning within a session ^16,83^. It is also inconceivable that determining which synapses to update (the credit assignment problem^109,110^) within the deluge of synaptic connections to BCI neurons can take place within a few minutes. As such, BTSP offers a more plausible mechanism that uses sensory feedback from the agent’s current state to ‘tag’ the synapses to BCI neurons that caused that state earlier in time, thereby permitting temporally specific information to act on local synapses deemed critical to potentiate (or depress)^111^.

We further provided support for our model by investigating both short- and long-term changes in firing characteristics in an exogenous BCI experiment. We first demonstrated that animals might not be able to learn rapidly with *any* ensemble, consistent with published reports ^16^. We then showed that the animal’s performance within the first few trials is a good predictor of their performance later in the trial and in subsequent sessions. These observations were supported by the ability of the animal to adopt a strategy based on how single neurons’ excitability could be altered with extended training. To preserve the E/I balance, synaptic scaling must slowly take place at the network level which likely involves non-BCI neurons to forge new ‘attractor’ states that can be rapidly attained with repeated practice, consistent with published work ^17^.

Our study nevertheless has important limitations. First, we used photo stimulation to maintain single neurons at activity levels *above* their baseline level. A topic for future work would be to design the experiment to also photo inhibit neurons *below* their baseline level that would mimic participation in decoding rules requiring differential modulation^112–114^, rather than pseudo integration, of individual neurons’ activities, such as the rule we used here in the optical clamp. Second, we characterized BTSP and IP in a small sample of the excitatory neurons class in V1 ^5,115^. We only tested the extent to which the clamp protocol causes neuronal adaptation to repeated inputs through habituation (response attenuation) but not sensitization (response enhancement). It would be important to extend the current work to larger sample size and other neuronal cell types, e.g. inhibitory cells, that are known to be sensitized to repeated inputs^71^, as well as in other brain areas known to be primary sources of BCI signals (e.g. motor cortex).

While the specific trends observed in implantable BCI experiments have been documented using mostly motoric, the mechanisms discussed extend beyond motor systems, for e.g., hippocampus ^44^. We therefore believe that our work is the first to propose a meta plasticity model that links rapid learning, representational drift, and BCI skill consolidation. Because of the causal nature of the transformation from neuronal activity to control signals that is unique to BCIs, more insights into how these disparate mechanisms could be working in synergy to facilitate learning and skill consolidation could be gained from this design.

In summary, our findings are critical to understand current limitations of BCI technology that impedes its progress and deployment in clinical and consumer applications. We propose that it an important step towards full autonomy of BCIs in everyday use would be to use neuroplasticity models like the one we propose here, combined with models of anticipated BCI use by the subject, to develop large scale, AI-powered brain-behavior BCI models that inform how decoders should *self-adapt* outside of actual BCI use. They would then be deployed regularly for use by the subject who might arguably find it less cognitively demanding to control and streamline its use in daily living.

## Acknowledgment

This study was supported by NINDS grant # NS93909. We thank the anonymous reviewers for their helpful comments.

The research has been conducted following NIH guidelines for research involving vertebrate animals under University of Florida’s approved IACUC Protocol # 202009788

## Supplementary Figures

**Supplementary Figure 1.**
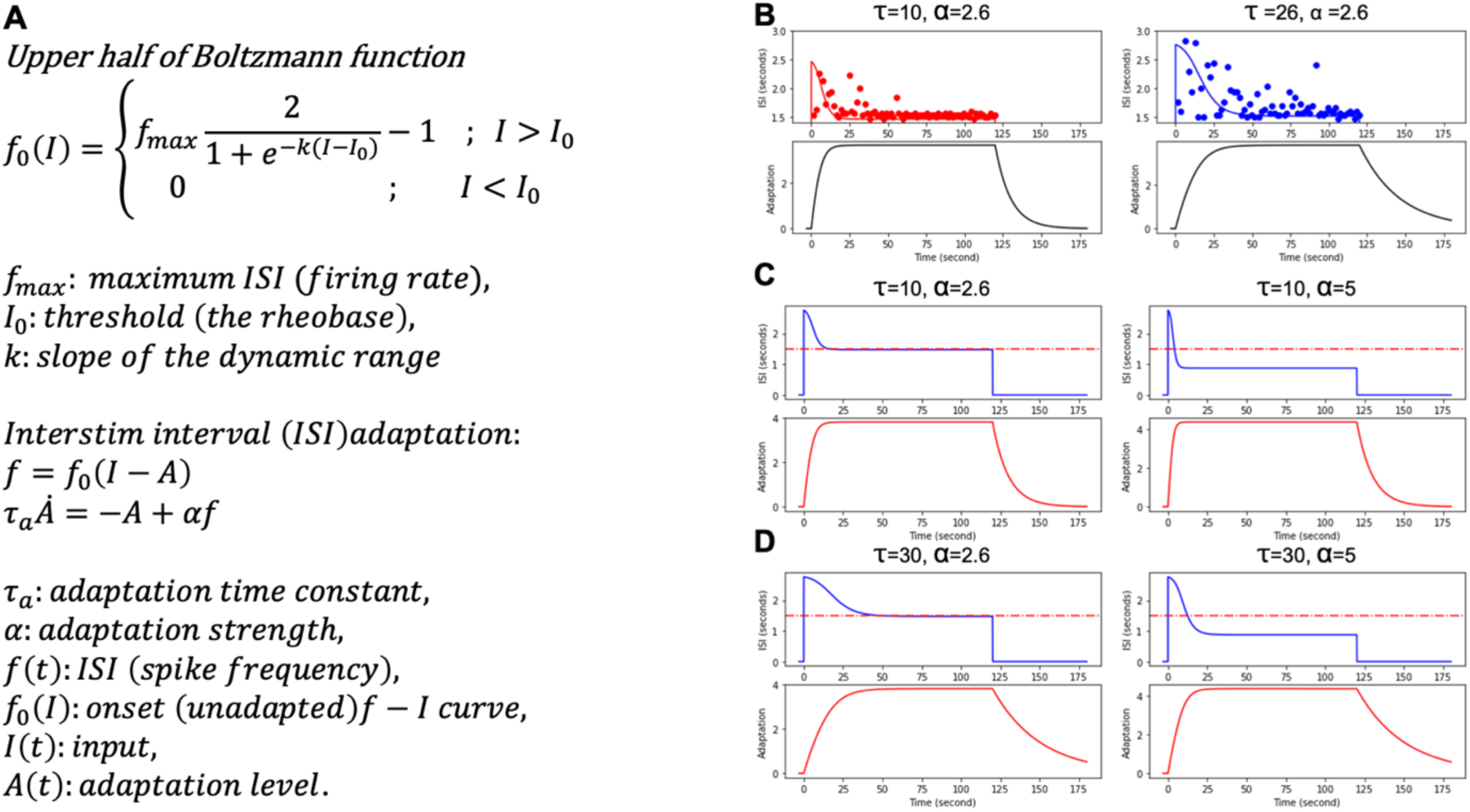
Model of Inter spike interval (ISI) adaptation as a function of clamp level. **A.** Boltzmann model equations describing how neurons’ ISI adapts to different inputs ^74^. **B.** Model prediction and actual data at different clamp thresholds (left 5 std, right 3 std) with adaptation time constant τ_a_ equal to 10 sec and 26 sec, respectively, and fixed adaptation strength *α* = 2.6 **C.** and **D**. ISI distribution generated by the model for different combinations of τ_a_ = 10 and 30 and α =2.6 and 5 that would hypothetically correspond to different input patterns.

**Supplementary figure 2.**
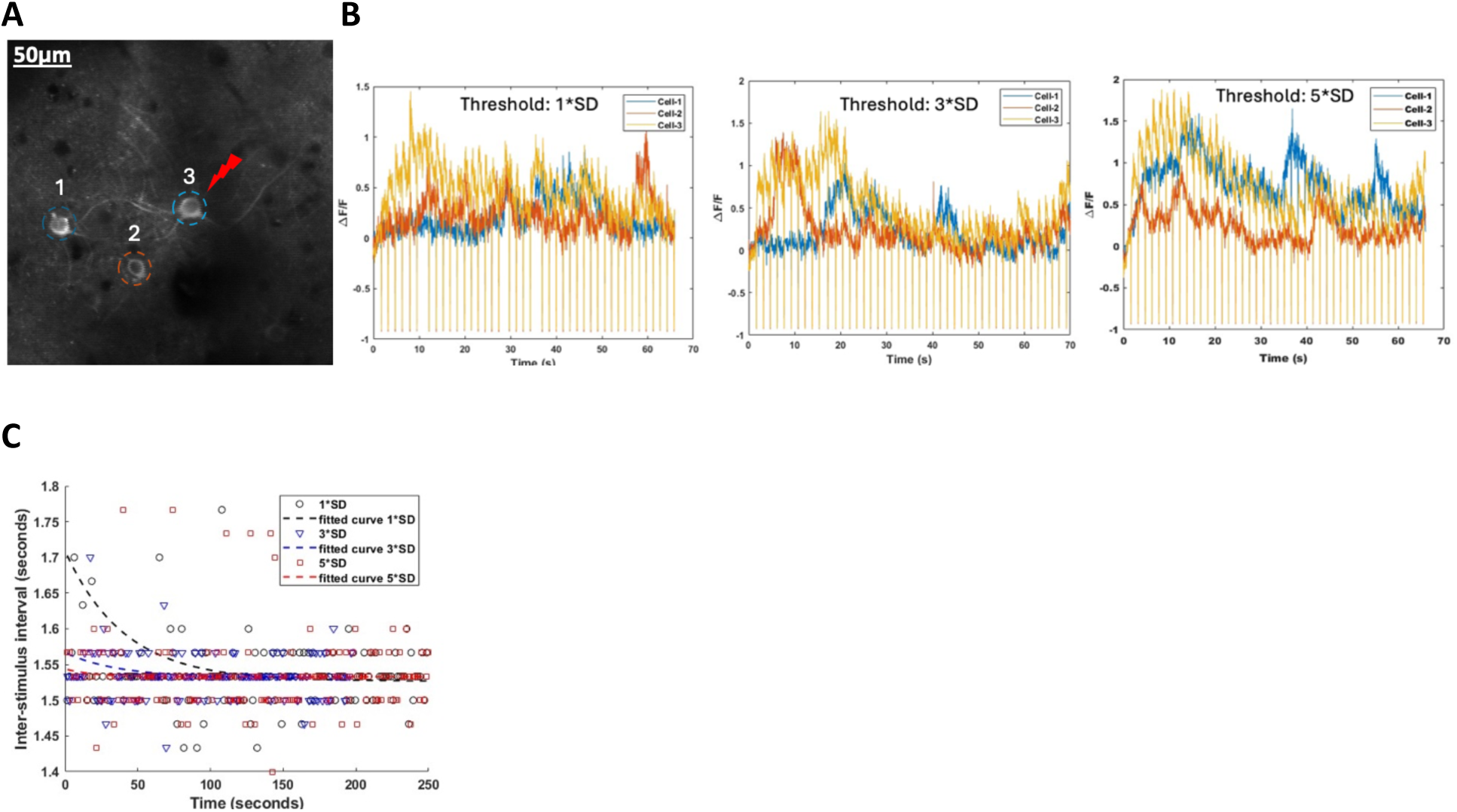
**A.** Effects of clamp on functional connectivity within local circuit. Sample FOV with 3 cells where cell 3 is chosen for closed loop stimulation at 1*SD, 3*SD and 5*SD. **B.** Raw traces of Ca^2+^ fluorescence from all 3 cells in **A**. **C.** ISI for cell 3 corresponding to the data in B, demonstrating different decay time constants as a function of target level (1*SD corresponds to 40.443 sec, 3*SD corresponds to 36.773 sec and 5*SD correspond to 19.332 sec).

**Supplementary Figure 3.**
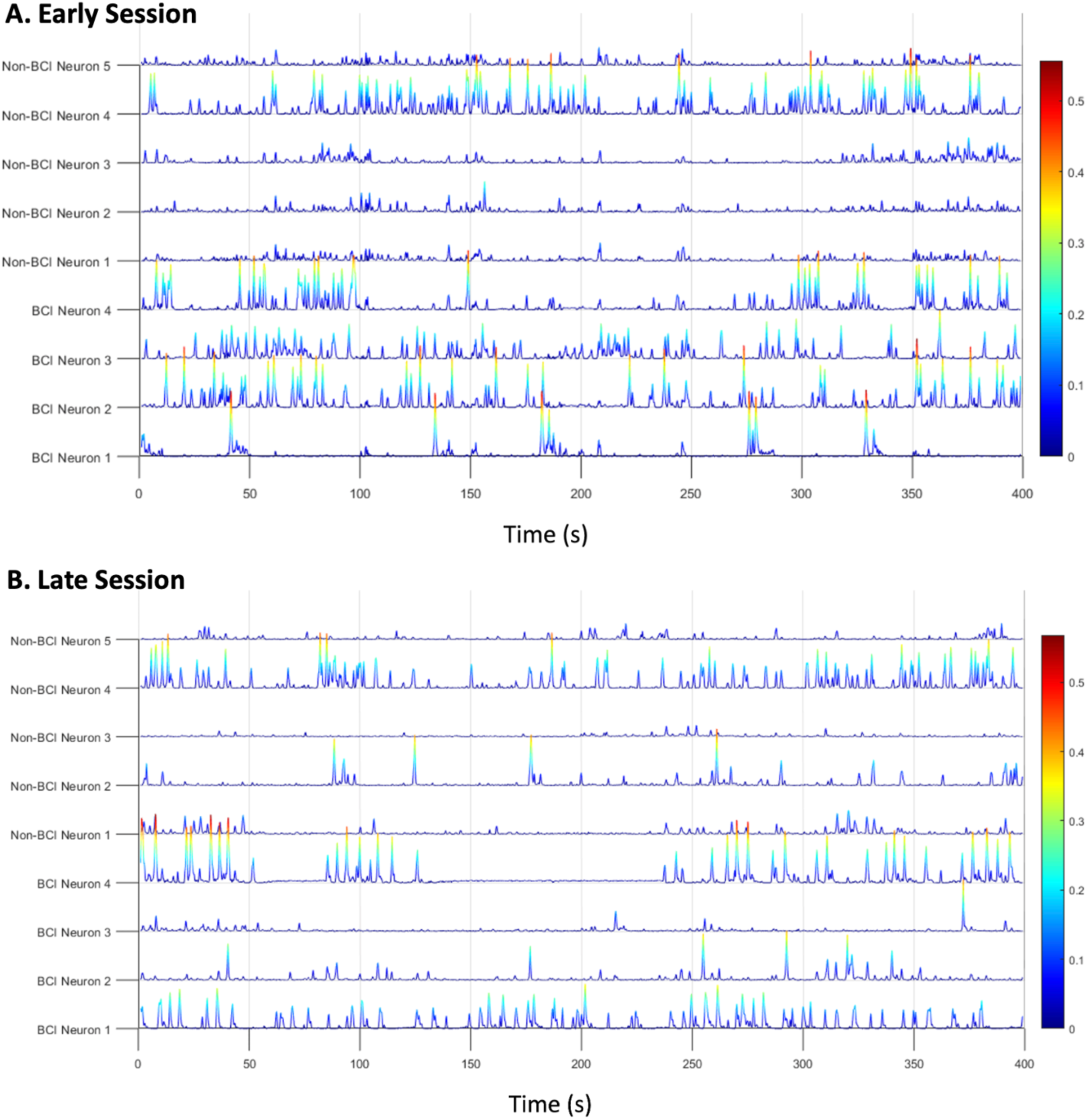
Spiking probability for E4 BCI neurons and non-BCI neurons during: **A.** the first BCI learning session and **B.** Session 13 taking place 2 weeks later.

**Supplementary Figure 4.**
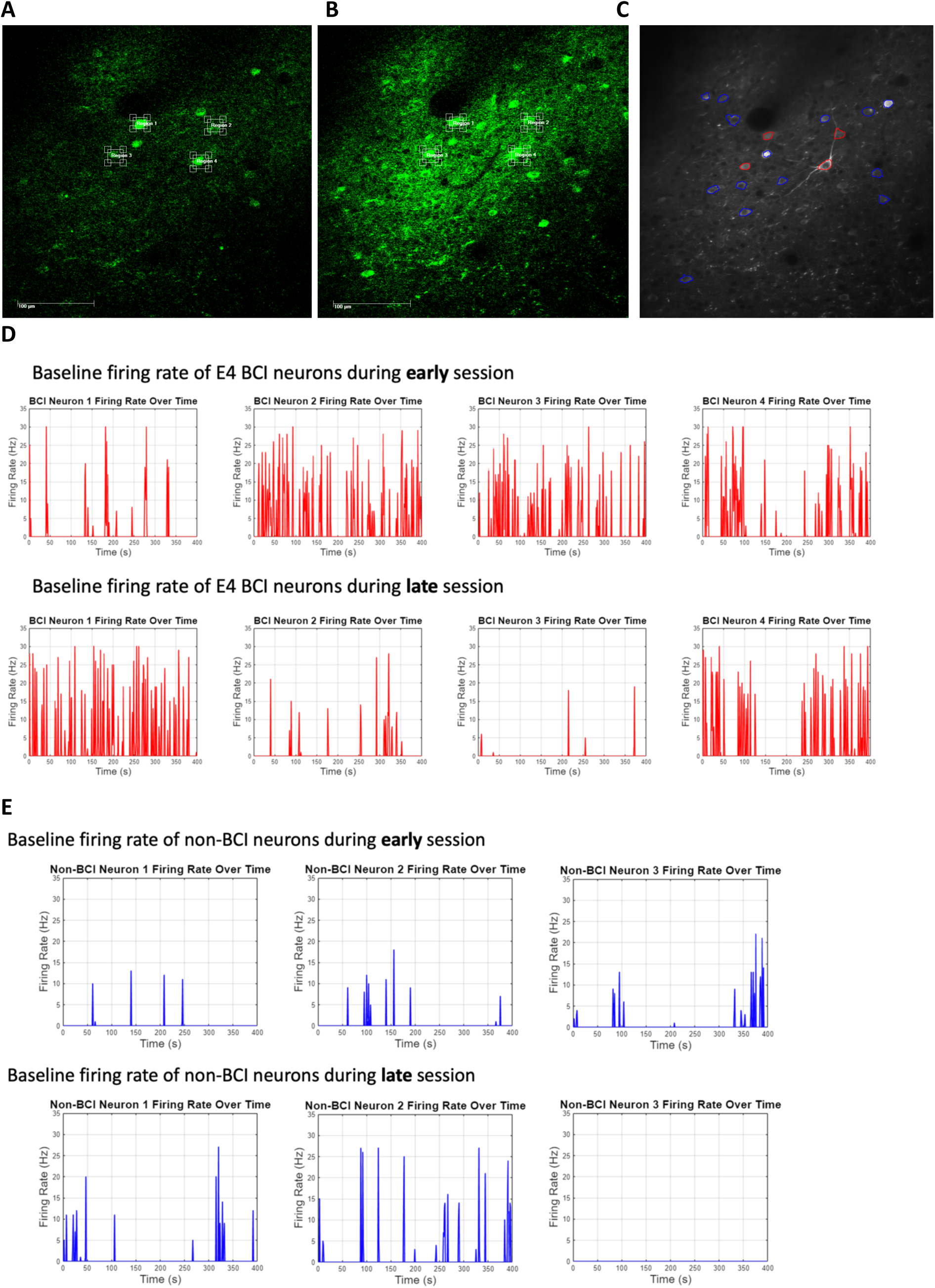
**A.** FOV used for the BCI experiment during the early and **B**. late session (2 weeks later) **C**. E4 ROIs consisting of the BCI neurons (blue) used throughout, Red ROIs are non-BCI neurons **D.** Baseline firing rate for 4 BCI neurons and **E.** three non-BCI neurons over 400 sec outside of BCI behavior.

## Supplementary Methods

### BCI experimental design

Water restricted mice expressing GCaMP7s in excitatory neurons were trained to volitionally modulate the activity of multiple ensembles of neurons for water rewards. Given an ensemble, the animal was rewarded for modulating the firing of a positive target pair (N+) above that of a negative target pair (N-) beyond a set threshold T1 according to the formula

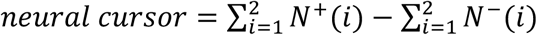

based on a decoder filter vector = [1, 1, −1, −1]. Lack of modulation within a maximum trial length of 30 sec indicated a failed trial, while inability to attain the target within this interval indicated an incomplete trial. Experiments were carried out over a number of sessions spanning multiple consecutive and non-consecutive days in which we alternated between ensembles that were either disjoint, partially or fully overlapping to quantify learning both within as well as across sessions.

The cursor feedback was encoded in the form of drifting gratings that increase in frequency with shorter distance towards the target and decrease with longer distance away from the target (supplementary video A). The animals are thus informed that they hit the target when the screen eventually becomes ‘all black’ and reward is dispensed, or that they had missed it when the screen eventually becomes ‘all white’ signaling a failed trial followed by a time out (4-sec). Inter trial intervals was indicated using a gray screen. A new trial starts by placing the cursor at an arbitrary ‘location’ in the virtual 1D maze.

### Supplementary video A

Illustration of a shifting drifting gratings feedback using an example summation of the modulated ensemble as a single “region of interest” within a square at bottom right corner illustrating the causal link between the fluorescence intensity change within the region when the cursor surpasses T1 and the corresponding movement of the drifting gratings in real time. When the cursor is below T1, a small noise component was used to move the gratings very slowly.

### Supplementary video B

Video illustrating the experimental setup in one session where the animal performed a number of successful consecutive trials in which the gratings started at a random initial position relative to the target (all black screen).

### Spiking Probability Inference

We used the CASCADE algorithm ^116^ to infer spiking probabilities of both the BCI and non-BCI neurons. CASCADE uses convolutional layers to extract local features of calcium signals, such as fast rise and slow decay, and merges them with other sequential modules (e.g., Long Short-Term Memory (LSTMs) to better capture long-term dependencies. We used the pre-trained Global EXC model as it exhibits remarkable generalization to many different neuron types and imaging conditions. This allowed us to apply robust inference of spike probabilities for the normalized calcium activity and to calculate the firing rate of each BCI and non-BCI neuron over time. When calculating the firing rate over time, we set the threshold of spike probability to 0.1 to suppress the background noise. If the spike probability value is less than 0.1, the firing rate at that time point would be set to 0.

